# Genomic characterization of three novel *Pseudomonas* phages within the subfamily of *Autographivirinae* isolated from organic waste

**DOI:** 10.1101/2020.01.20.910588

**Authors:** Jacob B Jørgensen, Amaru M Djuurhus, Alexander B. Carstens, Witold Kot, Cindy E. Morris, Lars H. Hansen

**Affiliations:** Department of Plant and Environmental Science, University of Copenhagen, Thorvaldsensvej 40, 1871 Frederiksberg, Denmark; INRAE, Pathologie Végétale, F-84140, Montfavet, France

## Abstract

Three phages targeting *Pseudomonas syringae* GAW0113 have been isolated from organic waste samples: Pseudomonas phage Bertil, Misse and Strit. The phages have double-stranded DNA genomes ranging from 41342 to 41374 bp in size comprising 50 to 51 open reading frames. The three phages genomes are highly similar and genomic comparison analyses shows that they all belong to the *Autographivirinae* subfamily of the family *Podoviridae.* The phages are however only distantly related to other members of this family, and have limited gene synteny with type-phages of other genera within *Autographivirinae*, suggesting that the newly isolated phages could represent a new genus.

## Introduction

*Pseudomonas syringae* is a gram-negative, rod-shaped bacterium. Strains of *P. syringae* are ubiquitous epiphytic plant pathogens and different strains can infect a wide range of important agricultural plant species, causing diseases such as bacterial canker of wild cherries[1] and kiwifruit[2], bacterial speck of tomato[3] and leaf blight in wheat[4]. The widespread problem with *P. syringae* in agricultural plants calls for an inexpensive, sustainable method for preventing or treating bacterial infections. Bacteriophage therapy is a promising approach to disease management[5], but requires a extensive library of relevant phages and understanding of phage-host interaction between phages and the *P. syringae* species. Here we present three phages isolated against *P. syringae* GAW0113[6]: Pseudomonas phage Bertil, Pseudomonas phage Misse and Pseudomonas phage Strit.

## Materials and methods

Samples of organic waste (OW) was collected, aliquoted and filtered with 0.45 μm syringe filters. The filtrate were tested for phage activity against *P. syringae* GAW0113 using a double agar overlay assay[7]. Plaques with distinct appearance or size were purified thrice. The phage DNA was extracted from lysate, as previously described[7] Phage DNA libraries were prepared using NEBNext Ultra II FS library Prep and Kit for Illumina (New England Biolabs, Ipswich, USA). The sequencing was performed on the Illumina iSeq platform as part of a flowcell (2 x 151, 300 cycles; Illumina). The genomes were trimmed and assembled in CLC Genomics Workbench version 12.0.3. Blastn[8] were used to search for sequence similarity against the non-redundant NCBI databases. Three of the phages are presented here because of low similarity with other known phages, but high similarity with each other.

The position of the terminal repeats were identified by the presence of two distinct spikes in read-start coverage at two nearby sites in the genome, encompassing areas with approximately double read-coverage. Open reading frames (ORFs) were identified automatically with GeneMark[9] and Glimmer[10] and revised manually in DNA Master[11]. The genomes were searched for tRNA sequences using Aragorn[12] and tRNAscan-SE[13], but none were found. The ORFs were manually annotated using both HHPred[14] against three downloaded databases the pFam 32.0, SCOP70 1.75 and PDB70 databases and blastp against the NCBI databases[8]. The nucleotide similarity heatmap between *Autographivirinae* phages was made with Gegenees version 3.1.0 (500 nt fragment size and 500 nt step size)[15]. A genome comparison between the isolated phages was made with Easyfig version 2.2.2. (default blastn settings with 2.9.0+ NCBI blast)[16].

## Results and Discussion

The genomes of the three phages are all double-stranded DNA, with sizes of 41373, 41342 and 41374 bp’s for Bertil, Misse and Strit respectively. All genomes contain terminal repeats of sligthly varying length (307 bp, 302 bp and 305 bp for Bertil, Misse and Strit respectively). The GC-content is 58.08 % for Bertil and Strit and 58.07 % for Misse. Bertil and Strit are almost identical except nine single nucleotide mutations and one deletion including a mis-sense mutation in *DNA polymerase I*. In total the phage genomes contains either 50 or 51 ORFs, all on the same strand. Misse lacks a 195 bp ORF situated in the middle of the genomes of Bertil and Strit, but contains a additional 100 bp non-coding region (fig. 1). The same 13 ORFs could be annotated with Hhpred (fig. 1) for each phage. These genes are mostly structural or involved in phage packaging and replication (fig. 1). No integrases or any other signs of lysogony where found in any of the three phages.

**Figure 1:**
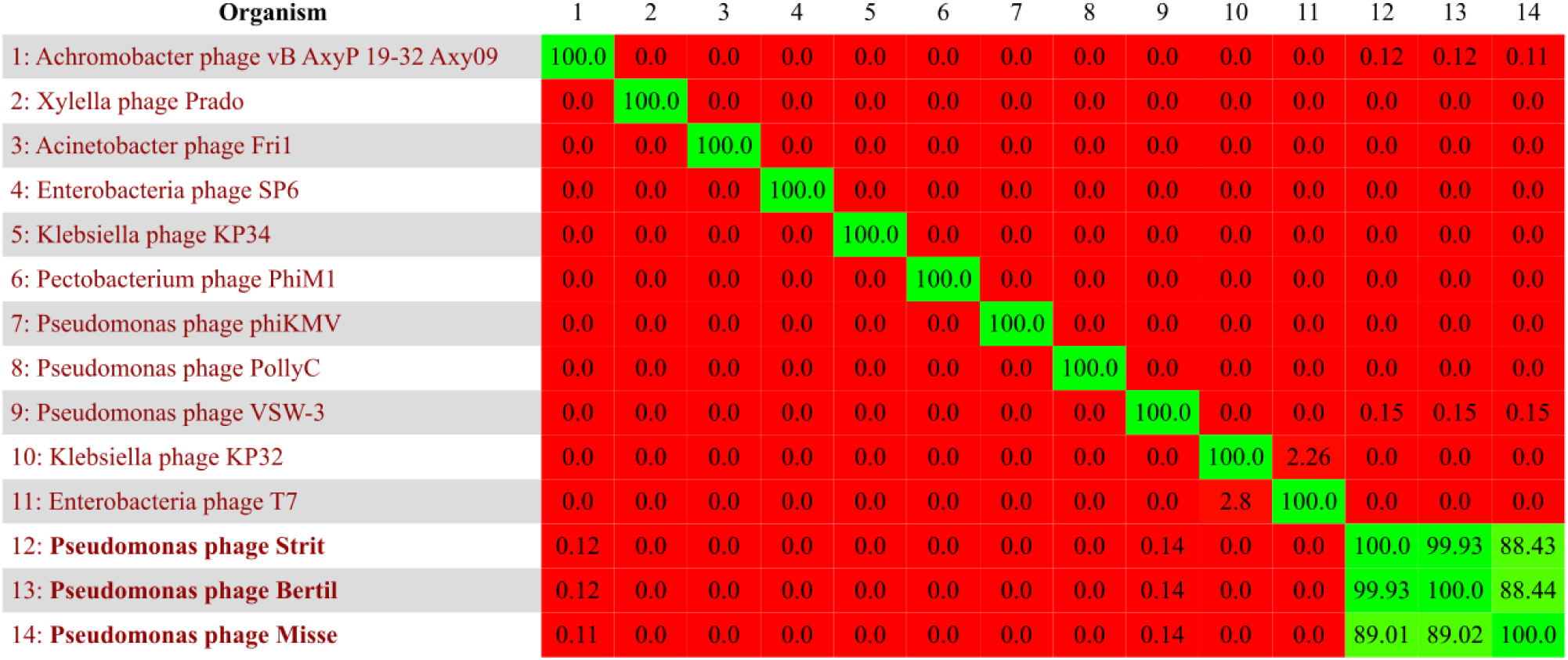
Heatmap showing all vs. all pair-wise nucleotide similarity between the type phages for the nine genera of the *Autographivirinae* subfamily and the three recently isolated *P. syringae* phages (in bold). The numbers and colors indicate similarity between the phage genomes from none or low (red) to high (green).

NCBI blastn shows that the three genomes have the largest similarity with *Achromobacter phage vB AxyP 19-32 Axy09* of the *Podoviridae* family and *Autographivirinae* subfamily (18 % Query cover and 70.1 % identity). The blastn showed 102 hits against phages in the *Podoviridae* family with 99 of the hits against phages in the *Autographivirinae* subfamily. Although there is little sequence similarity between the newly isolated phages and the type phages of *Autographivirinae* on a nucleotide level (fig. 2), gene synteny and gene size seems to be conserved for some of the structural genes (fig. 1). The similarity in gene synteny is however limited and is notably not the case for ex. the *DNA polymerase, terminase* and the *internal virion protein A* genes (fig. 1). Altogether, this suggests that the three phages could represent a new genus within the *Autographivirinae* subfamily.

**Figure 2:**
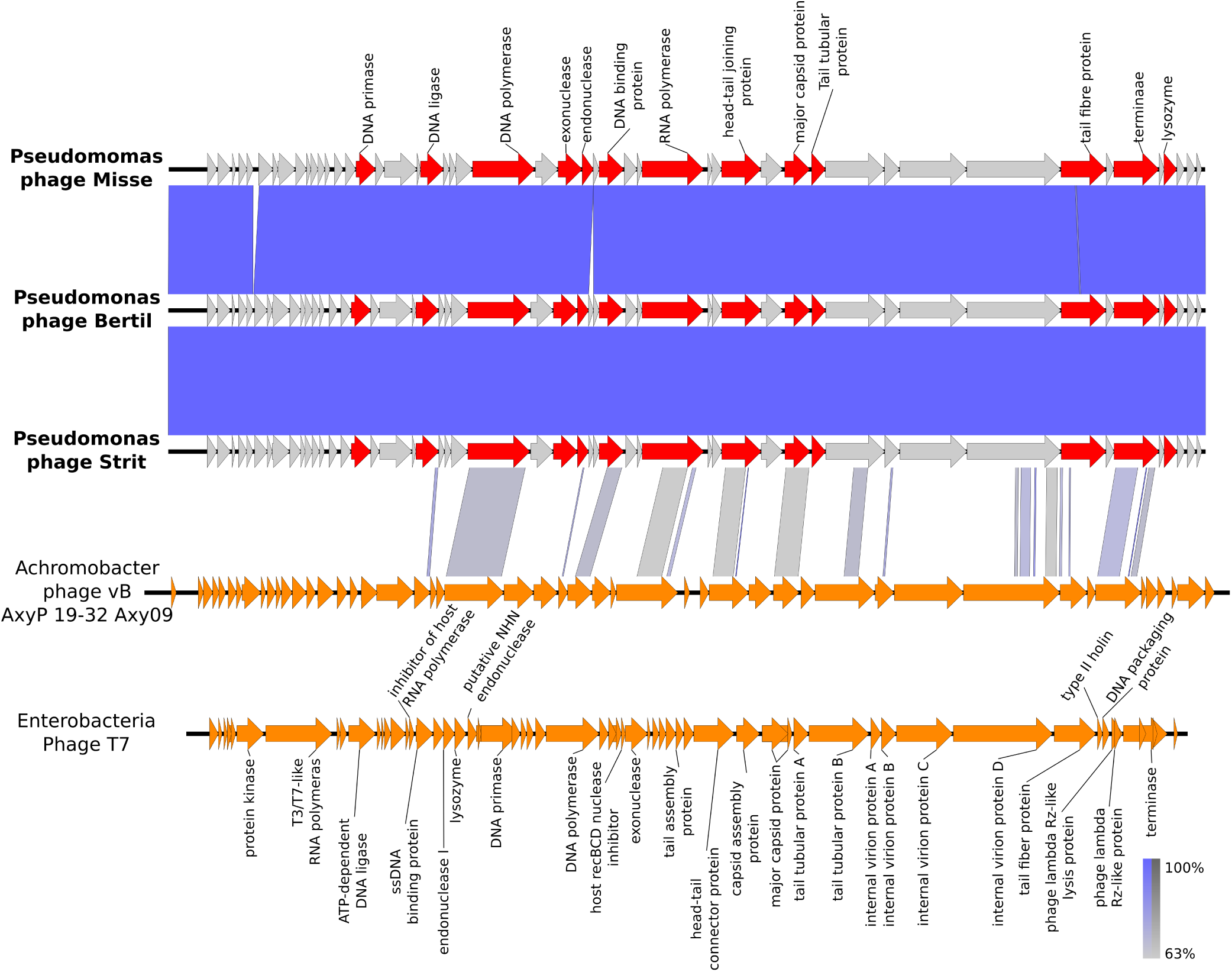
Comparison between Pseudomonas phages Bertil, Misse and Strit and 2 *Autographivirinae* phages with easyfig software (default blast settings). ORFs are represented by arrows indicating the reading direction. Pairwise nucleotide similarity between the different genomes is shown by color (from grey to blue). The annotated genes for Bertil, Misse and Strit are shown above the genome of Misse. Annotated genes are shown in red and hyp<u>o</u>thetical proteins are shown in grey.

The phages also share little identity with other known phages targeting *P. syringae*. Increased diversity within the known phages targeting *P. syringae* strains allow for development of more complex phage cocktails, which could decrease the possibility for the target bacteria to develop resistance to the treatment without losing fitness or virulence[17].

## References

[1] M. Scortichini, M. Biocca, and M. P. Rossi, “Pseudomonas syringae pv Morsprunorum on wild cherry for timber production: Outbreak and field susceptibility,” Eur. J. For. Pathol., vol. 25, no. 6–7, pp. 343–350, Dec. 1995.

[2] T. Y, S. S, I. T, T. S, and G. M, “Pseudomonas-Syringae Pathovar Actinidiae New Pathovar the Causal Bacterium of Canker of Kiwifruit in Japan,” Ann. Phytopathol. Soc. Jpn., vol. 55, no. 4, pp. 437–444, 1989.

[3] F. Sahin, “Severe outbreak of bacterial speck, caused by Pseudomonas syringae pv. tomato, on field-grown tomatoes in the eastern Anatolia region of Turkey,” Plant Pathol., vol. 50, no. 6, pp. 799–799, Dec. 2001, doi: 10.1046/j.1365-3059.2001.00622.x.

[4] J. Otta, “Wheat Leaf Necrosis Incited by Pseudomonas-Syringae,” Phytopathology, vol. 62, no. 10, pp. 1110–1110, 1972.

[5] M. Zaczek, B. Weber-Dabrowska, and A. Górski, “Phages in the global fruit and vegetable industry,” J. Appl. Microbiol., vol. 118, no. 3, pp. 537–556, 2015, doi: 10.1111/jam.12700.

[6] O. Berge et al., “A user’s guide to a data base of the diversity of Pseudomonas syringae and its application to classifying strains in this phylogenetic complex,” PLoS ONE, vol. 9, no. 9, 2014, doi: 10.1371/journal.pone.0105547.

[7] A. B. Carstens, A. M. Djurhuus, W. Kot, and L. H. Hansen, “A novel six-phage cocktail reduces Pectobacterium atrosepticum soft rot infection in potato tubers under simulated storage conditions,” FEMS Microbiol. Lett., vol. 366, no. 9, May 2019, doi: 10.1093/femsle/fnz101.

[8] S. Altschul et al., “Gapped blast and psi-blast: a new generation of protein database search programs,” FASEB J., vol. 12, no. 8, pp. 3389–3402, 1998, doi: 10.1007/s10535-010-0038-7.

[9] J. Besemer, A. Lomsadze, and M. Borodovsky, “GeneMarkS: a self-training method for prediction of gene starts in microbial genomes. Implications for finding sequence motifs in regulatory regions.,” Nucleic Acids Res., vol. 29, no. 12, pp. 2607–2618, Jun. 2001, doi: 10.1093/nar/29.12.2607.

[10] A. L. Delcher, K. A. Bratke, E. C. Powers, and S. L. Salzberg, “Identifying bacterial genes and endosymbiont DNA with Glimmer,” Bioinformatics, vol. 23, no. 6, pp. 673–679, 2007, doi: 10.1093/bioinformatics/btm009.

[11] DNA Master. Available: http://cobamide2.bio.pitt.edu/. [Accessed: 05-Dec-2019].

[12] D. Laslett and B. Canback, “ARAGORN, a program to detect tRNA genes and tmRNA genes in nucleotide sequences,” Nucleic Acids Res., vol. 32, no. 1, pp. 11–16, 2004, doi: 10.1093/nar/gkh152.

[13] P. P. Chan and T. M. Lowe, “tRNAscan-SE: Searching for tRNA Genes in Genomic Sequences,” in Gene Prediction: Methods and Protocols, M. Kollmar, Ed. New York, NY: Springer, 2019, pp. 1–14.

[14] J. Söding, A. Biegert, and A. N. Lupas, “The HHpred interactive server for protein homology detection and structure prediction,” Nucleic Acids Res., vol. 33, no. suppl_2, pp. W244–W248, Jul. 2005, doi: 10.1093/nar/gki408.

[15] J. Ågren, A. Sundström, T. Håfström, and B. Segerman, “Gegenees: Fragmented alignment of multiple genomes for determining phylogenomic distances and genetic signatures unique for specified target groups,” PLoS ONE, vol. 7, no. 6, 2012, doi: 10.1371/journal.pone.0039107.

[16] M. J. Sullivan, N. K. Petty, and S. A. Beatson, “Easyfig: A genome comparison visualizer,” Bioinformatics, vol. 27, no. 7, pp. 1009–1010, 2011, doi: 10.1093/bioinformatics/btr039.

[17] A. Svircev, D. Roach, and A. Castle, “Framing the future with bacteriophages in agriculture,” Viruses, vol. 10, no. 5, pp. 1–13, 2018, doi: 10.3390/v10050218.

